# Estimation of Laminar BOLD Activation Profiles using Deconvolution with a Physiological Point Spread Function

**DOI:** 10.1101/2020.08.04.236190

**Authors:** Irati Markuerkiaga, José P. Marques, Tara E. Gallagher, David G. Norris

## Abstract

**Background:** The specificity of gradient echo (GE)-BOLD laminar fMRI activation profiles is degraded by intracortical veins that drain blood from lower to upper cortical layers, propagating activation signal in the same direction. This work describes an approach to obtain layer specific profiles by deconvolving the measured profiles with a physiological Point Spread Function (PSF).

**New Method:** It is shown that the PSF can be characterised by a TE-dependent peak to tail (p2t) value that is independent of cortical depth and can be estimated by simulation. An experimental estimation of individual p2t values and the sensitivity of the deconvolved profiles to variations in p2t is obtained using laminar data measured with a multi-echo 3D-FLASH sequence. These profiles are echo time dependent, but the underlying neuronal response is the same, allowing a data-based estimation of the PSF.

**Results:** The deconvolved profiles are highly similar to the gold-standard obtained from extremely high resolution 3D-EPI data, for a range of p2t values of 5-9, which covers both the empirically determined value (7.1) and the value obtained by simulation (6.3).

**Comparison with Existing Method(s):** Corrected profiles show a flatter shape across the cortex and a high level of similarity with the gold-standard, defined as a subset of profiles that are unaffected by intracortical veins.

**Conclusions:** We conclude that deconvolution is a robust approach for removing the effect of signal propagation through intracortical veins. This makes it possible to obtain profiles with high laminar specificity while benefitting from the higher sensitivity and efficiency of GE-BOLD sequences.

## INTRODUCTION

Within the iso-cortex the type and density of brain cells varies following a laminar pattern. Following the canonical model (Felleman and Van Essen, 1991) there are distinctive patterns of laminar connectivity, corresponding to feedforward, feedback, and lateral interactions. Therefore, detecting neuronal activity at the laminar level is of interest to understand the hierarchical relationship between different brain regions and better understand brain function.

Although some initial attempts have been made using magnetoencephalography (Bonaiuto et al., 2018; Troebinger et al., 2014), functional MRI (fMRI) is to date the most popular non-invasive *in-vivo* technique to measure brain function at sub-millimetre resolution and obtain cortical depth-resolved activation signals (Koopmans and Yacoub, 2019; Norris and Polimeni, 2019).

The most widely used method in fMRI measures the Gradient Echo (GE) based blood oxygenation level dependent (BOLD) signal. The advantage of this method is that it has a higher sensitivity to activation and a higher efficiency compared to the other MR methods available for fMRI. The disadvantage is its reduced spatial specificity due to the contribution from large veins (Boxerman et al., 1995). In layer-specific fMRI the spatial specificity of the measured BOLD profile is degraded by the contribution from emerging or intracortical veins. These are the veins that are oriented perpendicular to the pial surface and drain blood uni-directionally from lower to upper layers. Therefore, part of the activation signal in a lower layer will propagate or leak into upper layers downstream. This phenomenon will be referred to as the “inter-laminar leakage-problem” in this manuscript.

There are a number of MR methods that suffer less from large vein contamination. These were originally developed for use in standard-resolution fMRI, but have been applied in laminar fMRI studies too. One of these methods consists of combining the standard GE-BOLD method with a differential paradigm (Cheng et al., 2001; Kashyap et al., 2018; Menon et al., 1997; Polimeni et al., 2010a; Sanchez Panchuelo et al., 2015). The signal resulting from the subtraction of the condition of interest and the control condition is expected to have a much lower contribution from large veins. SE-based sequences represent another way of obtaining an activation signal with little contribution from large veins (Goense and Logothetis, 2006; Harel et al., 2006; Harmer et al., 2012; Kemper et al., 2015; Yacoub et al., 2007; Yacoub et al., 2009; Zhao et al., 2006). The 180° pulse refocuses the extravascular signal in the static dephasing regime, which is the source of the largest contribution in GE-BOLD at 3T and above (Cheng et al., 2015). The S2 signal of the unbalanced SSFP sequence is believed to be dominated by T2 contrast, therefore the BOLD signal is expected to have only a small contribution from larger veins at high-fields (Barth et al., 2010; Ehses et al., 2013; Goa et al., 2014). An important method for obtaining an activation signal without venous contribution, that can be applied in laminar studies, is the measurement of cerebral blood volume (CBV) changes using the vascular space occupancy (VASO) method (Finn et al., 2019; Guidi et al., 2016; Huber et al., 2015; Huber et al., 2017; Lu et al., 2003). The activation signal measured with this method is proportional to the change in vascular blood volume. As the largest portion of blood volume change is located in the arterioles and capillary bed, (Krieger et al., 2012) this technique is not weighted towards larger post-capillary vessels.

The alternative methods to GE-BOLD mentioned above present different specific technical difficulties of their own (for example: sensitivity to motion, risk of exceeding energy deposition limits at high static magnetic field strengths, inefficient tagging or inversion of the blood signal). Nonetheless, they all have in common a lower sensitivity to activation and temporal resolution compared to the standard GE-BOLD based fMRI approach.

This manuscript examines a method to deal with the inter-laminar leakage problem in the steady state. It consists of deconvolving the measured GE-BOLD activation profile averaged across the cortex with the spatial physiological point spread function (PSF) of the BOLD response to laminar activation. This physiological PSF characterises the BOLD signal leakage from the layer of activation to layers downstream. As it is not generally possible to measure the PSF *in vivo*, the PSF estimated in Markuerkiaga et al. (2016) using a model of cortical vasculature is used. The strength of this method is that it is expected to deliver BOLD activation profiles with little inter-laminar leakage while allowing the use of data acquired with the simplest fMRI sequence that has the highest efficiency and sensitivity. The method presented in this manuscript and the spatial GLM approach (Kok et al., 2016; Polimeni et al., 2010b; van Mourik et al., 2019) do not address the same problem. The deconvolution approach aims at removing the effect of draining veins from the functional profiles whereas spatial GLM is a purely geometric approach that aims to reduce partial volume effects, and is not necessarily related to functional acquisitions.

The deconvolution approach has previously been proposed (Markuerkiaga and Norris, 2016) and subsequently adopted by other users (Marquardt et al., 2018). It is attractive because although deconvolution can be an unstable process, here it is equivalent to a subtraction of responses in lower layers from the responses in higher layers, which is mathematically more robust. By use of a more sophisticated modelling approach it can also be extended to the dynamic situation (Havlicek and Uludag, 2020). In the current contribution the validity of the deconvolution approach is assessed on previously published profiles acquired at 7T. First the approach is validated on the profiles obtained using an ultrahigh resolution 3D-EPI acquisition approach (Fracasso et al., 2018). The profiles obtained in this study were grouped into strong, middle and weak linear trends, whereby the latter were assumed to have a very small contribution from intracortical veins. These latter profiles are considered as the gold-standard of leakage-free activation profiles in this manuscript. Successful deconvolution of the full profile should hence yield a profile similar to those with little contribution from intracortical veins. We then deconvolve the profiles at different TEs obtained using a multi-echo 3D-FLASH sequence (Koopmans et al., 2011). In this dataset, BOLD profiles are echo dependent, mostly due to the strong TE-dependency of the intravascular venous contribution, but the underlying neuronal state is not. This feature is exploited to perform a data-based estimation of the PSF. Lastly, the sensitivity of the deconvolution results to the choice of parameters used for the PSF is studied.

## MATERIALS AND METHODS

### Mathematical description of the inter-laminar leakage

In the steady state, the measured activation profile can be understood as the weighted sum of the laminar BOLD responses convolved with the corresponding layer specific physiological point spread function (PSF). This can be expressed as *Y* = *X*\*β + ε, where *X* is a matrix of layer specific PSFs, β is the underlying laminar BOLD response, and ε is the error term. Specifically, the elements of the equation Y = X*β + ε are:

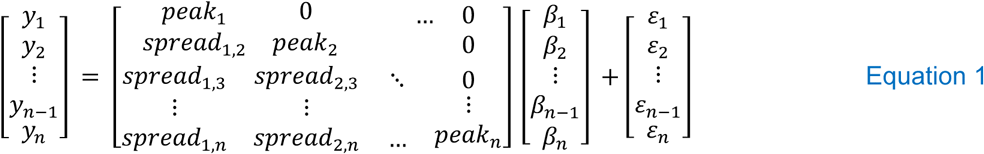

where *y*_*i*_ is the BOLD response measured at bin *i, peak*_*i*_ is the peak value of the PSF measured at bin *i* (BOLD response directly related to the local neuronal response), *spread*_*i,k*_ is the spread of the PSF measured at bin *k* that results from the activation upstream in bin *i. β* is the underlying laminar BOLD response free of inter-laminar leakage and *n* is the number of bins considered in the analysis. *Bin* indexes the sampling points across the cortex, and runs from 1, in the bin adjacent to the white matter boundary, to n, the bin at the grey matter/CSF boundary. ε is the error term that accounts for a combination of measurement noise and potential deviations of the model from the biophysical system. As the cortex is drained by intracortical veins unidirectionally from lower cortical layers towards the cortex, signal will only spread towards the cortical surface and the physiological PSF will be skewed in this direction. That is why, *X* is lower triangular matrix. If the layer specific Point Spread Function is known, i.e. the magnitude of the *peaks*_*i,j*_ and *spreads*_*i,j*_ in Equation 1, then the underlying laminar BOLD activation profile can be obtained by solving for β, as 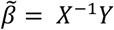.

### The physiological Point Spread Function

It is not possible to measure the physiological Point Spread Function (PSF) *in-vivo* in humans due to the difficulty of obtaining the response of a single layer in a region, without simultaneously generating a response in other layers within the region due to intra-regional laminar connections. A model of cortical vasculature was previously developed to estimate the physiological PSF (Markuerkiaga et al., 2016). In that work, a simple model of cortical vasculature based on published histological studies focusing on the primary visual cortex was developed. A previously-developed BOLD signal model (Uludag et al., 2009) was then applied to obtain the positive BOLD response across the cortex. As the simulated neuronal response, the features of the underlying vasculature and the resulting BOLD response were known, it was possible to estimate the physiological PSF. This model-based estimation of the physiological PSF will be used in this manuscript (in updated form) to obtain leakage-free activation profiles.

The physiological PSF of the laminar response in a small region of the cortex will depend on the baseline physiological conditions and the following two factors: the presence and size of emerging veins and the orientation of the cortex with respect to B_0_. The physiological PSF estimated in Markuerkiaga et al. (2016) applies to the situation in which the cortical profiles to be deconvolved are the result of integrating over a convoluted patch of cortex, and thus effectively averaging over all orientations with respect to B_0_. Baseline physiological conditions will affect the magnitude of the response, but not the shape. In addition, the estimated PSF in that work characterized the PSF for stimuli longer than 1s - when the cortex has reached a steady state after the stimulus onset (Markuerkiaga et al., 2016)

Figure 4b of Markuerkiaga et al. (2016) shows that the estimated PSF is very similar between layers. It shows a peak in the layer of activation and a rather constant tail towards the pial surface. Hence, the response can be approximated using two parameters, the peak at the site of activation and a constant tail from the site of activation to the pial surface. This implies that the ‘spread’ values in Equation 1 do not vary between bins, and hence that these can be replaced by a single ‘tail’ value for each column in matrix X of Equation 1, and furthermore that the ratio of the peak to the tail value is independent of the layer. In this scenario, matrix *X* in Equation 1 will only have two parameters and can be rewritten as shown in Equation 2.

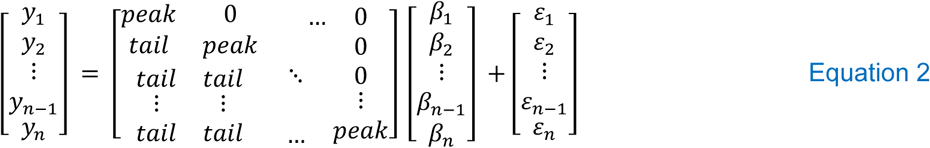

The peak and tail values might vary between subjects due to the inter-subject variability of the BOLD response. However, given that the vascular architecture is similar between individuals, their ratio (the peak to tail ratio, p2t) can be expected to show less variance. There are two ways of determining the elements of matrix *X* in Equation 2. One of the options, which we will refer to as normalised, is to set *peak* = *y*_*1*_ and tail = *y*_*1*_/p2t, in which case the underlying profile of the BOLD response, *β*, will be relative to the response in the lowest bin. The other option is to write the matrix as all ones on the diagonal and 1/p2t in the lower diagonal. In this second approach, the laminar BOLD profiles will not be normalised and differences in the average magnitude of the profiles between subjects would reflect differences in the average BOLD response between them, and variations in the profiles would represent both the underlying neuronal activity and differences in the underlying vascular density.

The consequences for the laminar profiles of: using an exact PSF for each layer; normalising this exact PSF to the response at the lowest bin; or replacing the normalised, layer-specific PSFs with a single normalised p2t value that is constant across the cortex, are described in the *Supporting Information (part 1)*. The results of this analysis show that: a deconvolution with the exact PSF corrects both for leakage and differences in vascular density; normalisation means that only leakage is corrected, and that there is little difference between using a true PSF form, or a single p2t-value for the entire cortex (see SI:Figure 1). The latter two approaches approximate the deconvolved profile from what could be obtained by a spin echo acquisition, which is sensitive to variations in vascular density.

**Figure 1:**
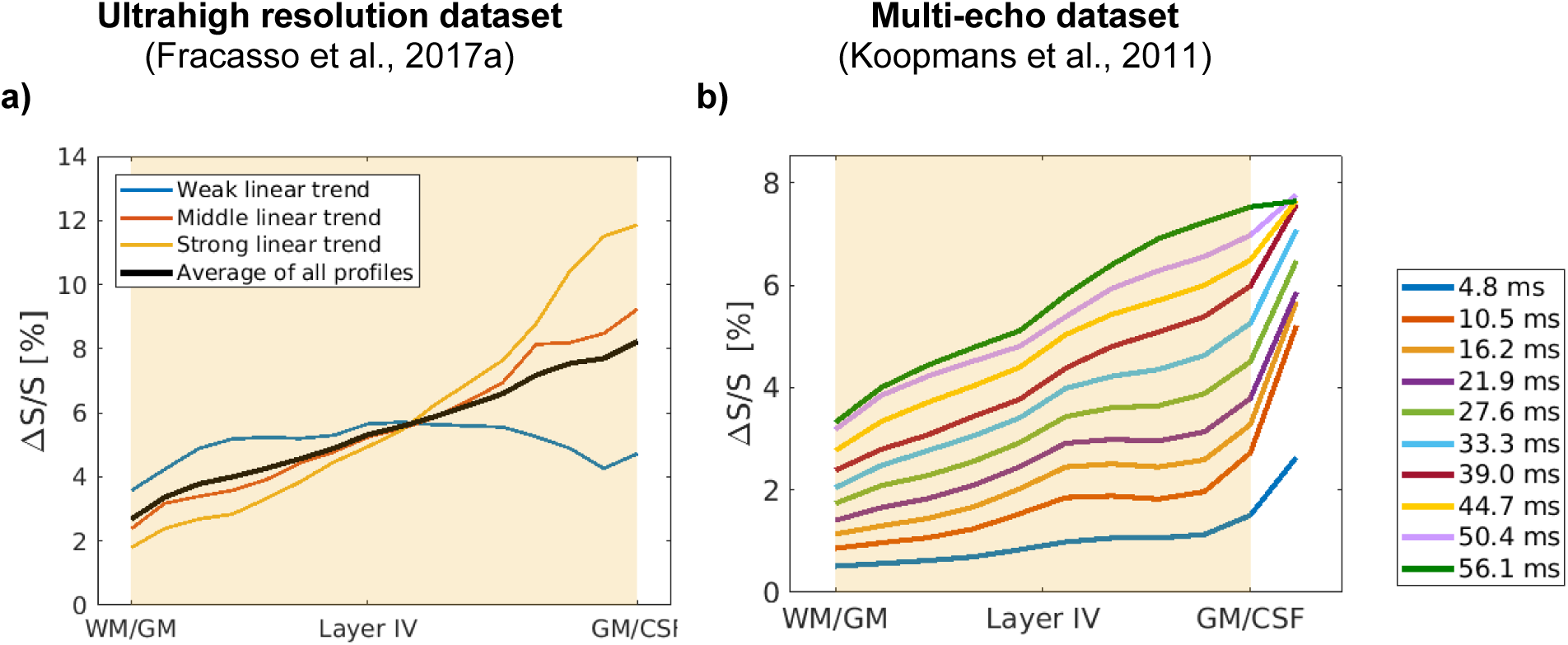
GE-BOLD cortical profiles used to test the deconvolution approach in this manuscript. The profiles shown are from an exemplary subject in each dataset. The data-points within the beige-shaded region were used for the deconvolution. The average of all profiles in the ultrahigh resolution dataset, a), was obtained using Equation 6.

#### Magnitude of the p2t and its adjustment for different cortical sampling densities

Following (Uludag et al., 2009), the parameters used for estimating the p2t at 7T in Markuerkiaga et al. (2016) assumed no intravascular venous contribution to the BOLD signal change at this field strength. However, there are a number of studies performed at 7T that indicate that there is an intravascular contribution to the BOLD signal at the echo times typically used in fMRI experiments (Blockley et al., 2008; Koopmans et al., 2011; Poser and Norris, 2009) and that this amounts to ∼ 8% of the total BOLD response at TE=T2*_GM_ at 7T. In order to have a more experimentally accurate p2t value, the parameters in Markuerkiaga et al. (2016) were updated so that the venous intravascular contribution at 7T would be ∼8% (see *Supporting Information*, part 2). The resulting p2t value at TE=T2*_GM_ at 7T is then 6.3, instead of the 5.1 previously obtained in the absence of intravascular contribution.

The p2t values presented in the previous paragraph assume that the cortex has been divided into 10 bins. If a different number of bins is used, then the p2t has to be adjusted according to the following formula:

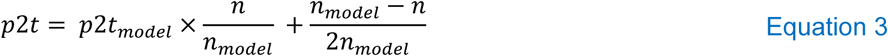

where *n* is the number of bins across the cortex in the acquired data and *n*_*model*_ is the number of bins used in the model used to estimate *p2t*_*model*_ (in this case, *n*_*model*_*=*10). See *Supporting Information part 3* for the derivation of Equation 3.

#### Data based estimation of the physiological PSF

It is not possible to measure the physiological PSF non-invasively *in-vivo*. However, assuming that it is valid to characterise the physiological PSF using a single value of the p2t ratio, then a multi-echo dataset can be employed to perform a data-based estimation of this parameter. In this manuscript we utilise a multi-echo FLASH acquisition, which gives an activation profile for each echo time for the same underlying neural activity. The response at each echo time is described by Equation 2. The parameters of the physiological PSF in matrix *X* will vary with TE (as described below) but the pattern followed by the underlying laminar response, β, will be TE-independent. Therefore, when activation profiles at different echo times are combined, the system will not be underdetermined and both the TE-dependent peak and tail magnitudes (and hence the p2t) and the TE-independent lamina BOLD responses can be estimated. The multi-echo 3D-FLASH sequence used had 10 echo times ranging from TE= 5.7 ms to TE= 56.1 ms (see next subsection for more details on these data). In order to better compare profiles acquired at different TEs it was found necessary to also model the signal contribution from the pial leading to a slight extension of Equation 2. This is particularly important for short TEs where the intravascular contribution at the pial surface can be large. Two parameters, the pial response, *pial*, and the partial volume factor, *α*, are introduced to account for partial volume effects with the pial surface signal as follows:

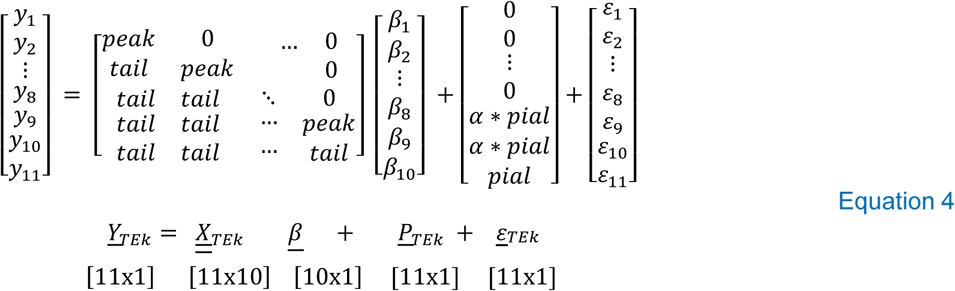

where *α* is the fraction of pial partial volume and can spread up to two bins (equivalent to one voxel) below the cortical surface (see Figure 3 in (Koopmans et al., 2011)). The parameter *pial* refers to the magnitude of the pial signal, i.e. *y*_11_, which is left for the algorithm to solve. All other parameters are as defined for Equation 2. Equation 5 shows how data for the different echo times, the elements in Equation 4, are combined to estimate p2t based on the experimental data. As mentioned earlier, the underlying neural- and hemodynamic responses do not vary between echo times, and the profile of the leakage free BOLD response, β, across the cortex will not vary either. The magnitude of the leakage-free profile is in principle TE-dependent, but as the peak and tail values are calculated for each echo time, any differences in the average magnitude of β across echo times will lead to a scaling of the corresponding peak and tail values.

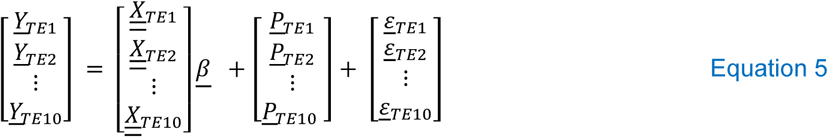

**Figure 2:**
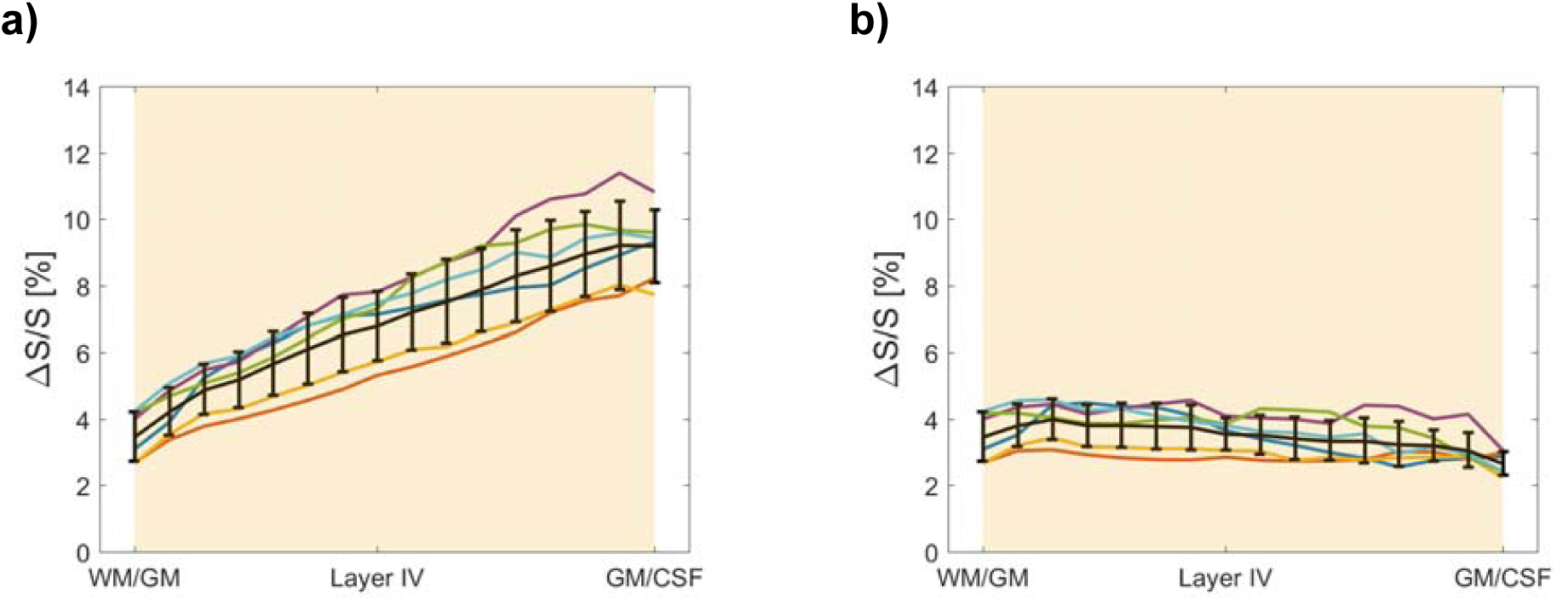
**a)** Weighted average profile of the weak, middle and strong linear profiles of the positive BOLD response in (Fracasso et al., 2017a). **b)** The corresponding deconvolved profiles. The different colours refer to different acquisitions, 5 subjects with one subject acquired twice. The black line shows the average ± std across subjects.

**Figure 3:**
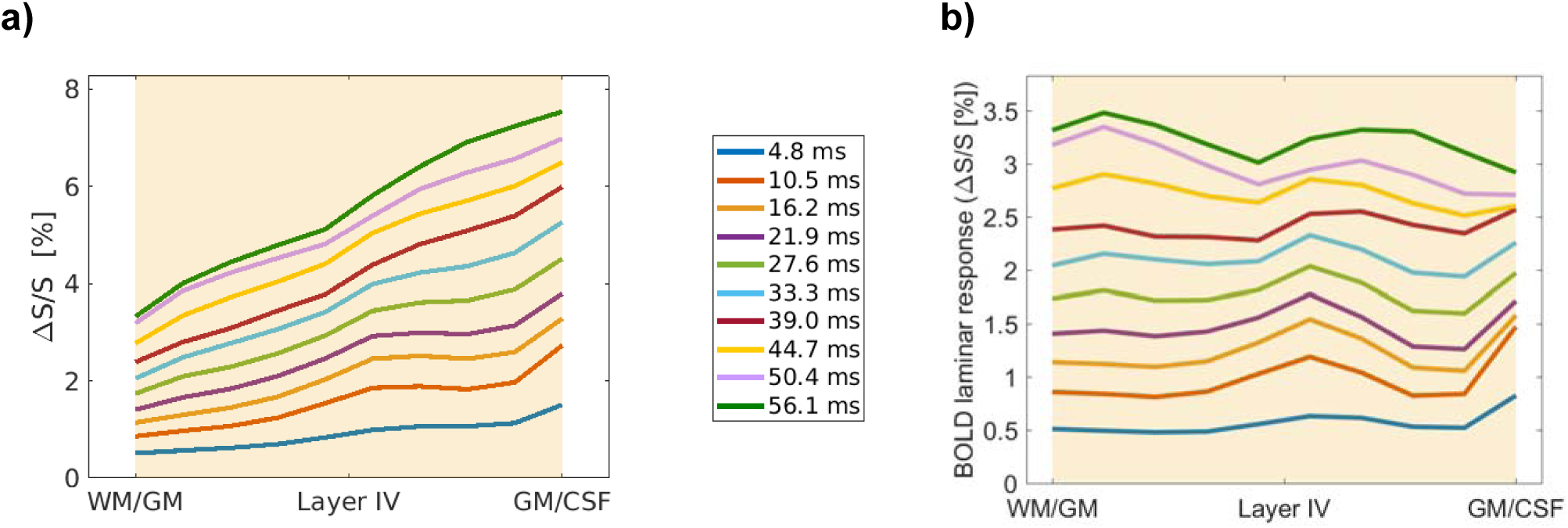
**a)** Measured cortical activation profiles averaged over subjects in the multi-echo dataset (Koopmans et al., 2011) as a function of TE. **b)** Deconvolved profiles, averaged over subjects, that result from deconvolving the activation profiles in a) with the corresponding, echo-time specific, p2t estimated using the vascular model.

In Equation 5 there are 41 parameters to estimate (10 underlying leakage-free BOLD responses β, one per cortical bin, one peak, one tail and one pial surface value per echo time, and one partial volume fraction) and 110 observations (11 cortical bins, including the pial surface, for each of the 10 echo times). Least squares non-linear data-fitting using the ‘Trust region algorithm’ as implemented in MATLAB 2017a (The MathWorks, Inc.) was used to approximate the solution of the overdetermined system. Apart from the description of the system to solve, this solver requires initial values of the parameters for the first iteration, as well as upper and lower boundaries. Lower boundaries have all been set to 0 and upper boundaries to 10. Initial parameter values were set in the following way:

- Leakage-free BOLD response, β: values were randomly picked at each cortical bin from a uniform distribution between 0 and 10.
- Peak values: randomly picked for each TE from a uniform distribution between 0 and 10.
- Tail values: the peak value divided by a plausible peak to tail ratio (p2t). This peak to tail ratio was randomly chosen from a uniform distribution between 4 and 8. This range was chosen based on the results presented in Figure 5, and because the p2t estimated in the model was 6.3.
- Pial values: randomly picked for each TE from a uniform distribution between 0 and 10.

**Figure 4:**
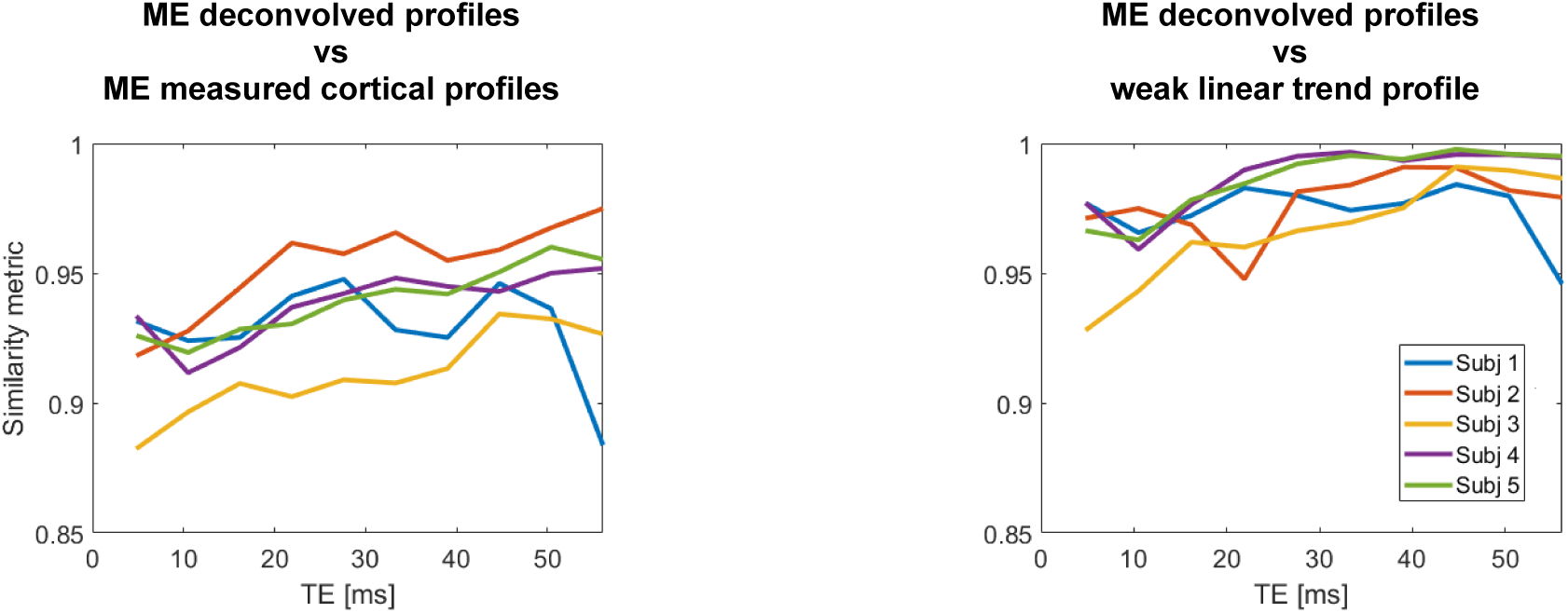
**Left:** similarity metric for the deconvolved profiles in the multi-echo dataset compared with their own (non-deconvolved) measured cortical profile. **Right:** similarity metric for the deconvolved profiles in the multi-echo dataset and the weak linear trend profile in the high-resolution dataset (blue line in Figure 2a). The colour legend is the same in both graphs. The similarity metrics are significantly different from each other between the graphs (p=0.027, α=0.05).

**Figure 5:**
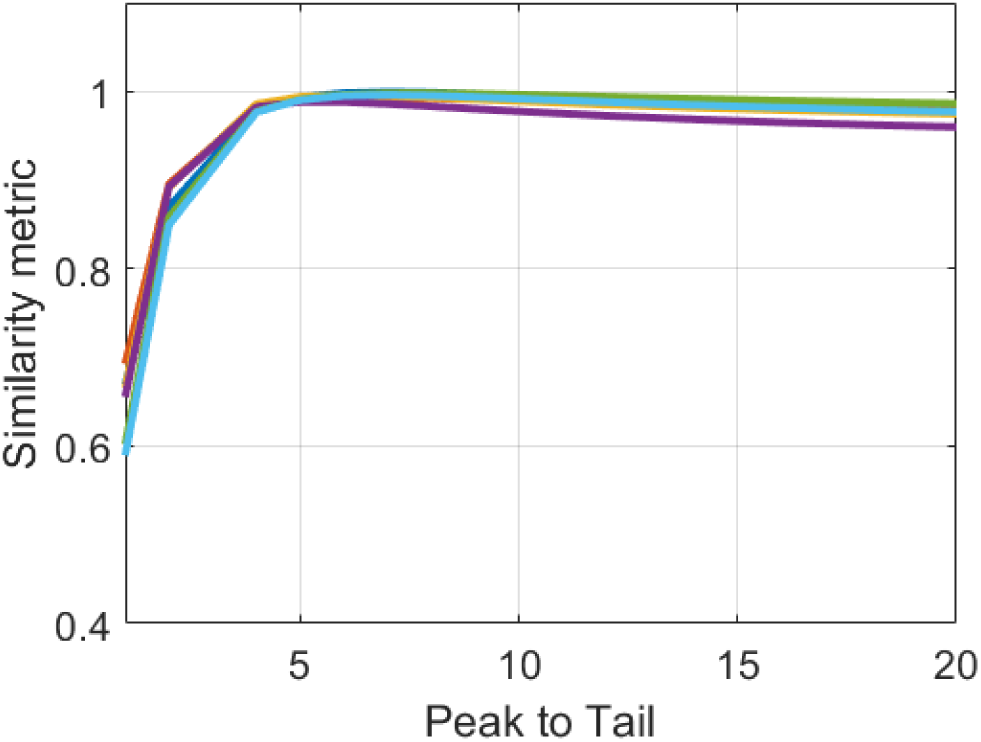
Similarity between the average cortical profile in the ultrahigh resolution dataset (shown in colour in Figure 2a) deconvolved with p2t values ranging from 1 to 20, and the corresponding weak linear profile from the same acquisition. The similarity index reaches its maximum value at about 5 for each subject, and then declines gently as the p2t increases further.

As no constraints on the profiles of the βs were set, very large and small values of p2t could also represent numerical solutions for the system, although they are not physiologically plausible. That is why a loose relationship following physiologically plausible values was set between the initial values of the peak and tail parameters (i.e. the third item in the list above). In order to avoid that the parameter estimations were strongly biased by the initial parameter values fed to the solver. The procedure was repeated 100 times and the median of the 100 p2t values calculated using this procedure was considered to be the p2t for each TE estimated by this method.

### Description of experimental datasets

The approach to obtain leakage-free BOLD laminar responses was tested in two functional datasets that were originally acquired for other purposes: an ultrahigh-resolution dataset with 0.55 mm isotropic voxel size (Fracasso et al., 2018) and a multi-echo FLASH dataset (Koopmans et al., 2011). A detailed description of the MR acquisition protocols and the extraction of the cortical activation profiles can be found in the original articles. Here, the most relevant parameters necessary to understand the nature of the profiles used for the deconvolution are given (see Table 1 for a summary).

**Table 1:** Relevant acquisition parameters of the experimental datasets used in this manuscript to test the deconvolution

| Parameters | Multi-echo dataset<br>(Koopmans et al., 2011) | Ultrahigh resolution dataset<br>(Fracasso et al., 2017a) |
| --- | --- | --- |
| Voxel size | 0.75 mm isotropic | 0.55 mm isotropic |
| No. bins: | Downsampled to 10 within GM | 16 within GM |
| Sequence | 3D-FLASH, multiecho | 3D-EPI |
| Vol TR/TR/TE | 90 s/66 ms/5.7-56 ms | 6.5 s/54 ms/27 ms |
| Echo train length | - | 53 ms |
| Flip angle | 15° | 20° |
| Visual stimulus: | Flickering checkerboard | Semi-field flickering checkerboard |
| No. subjects | 5 | 5 (one of which was scanned twice) |

#### Ultrahigh-resolution dataset

The ultrahigh-resolution dataset was acquired at 7T using the 3D-EPI sequence and a dedicated coil array positioned under the occipital lobe (Fracasso et al., 2018). The response to a checkerboard-like stimulus covering half of the visual field was measured in ipsilateral and contralateral V1, obtaining a negative and positive BOLD response, respectively. The voxel size was 0.55 mm isotropic. The cortical boundary was manually segmented and, following the equivolume principle, 16 surfaces across the cortex were generated. The 16 points of the corresponding node (or vertex) at each of the surfaces make up the sampling points for a given profile.

Following Fracasso et al. (2018), data were divided into profiles presenting a weak, middle or strong linear component (Figure 1a shows the profiles of the positive BOLD response for a sample subject).The argument for this parcellation was that at this resolution, some of the profiles will not contain large emerging veins and will therefore be dominated by the laminar response.

The first 40% of the profiles with the weakest linear trend were averaged to obtain the weak linear trend profile. The strongest 30% linear trend were averaged to obtain the strong linear trend and the remaining 30% generated an intermediate trend. As the p2t value estimated in Markuerkiaga et al. (2016) assumes that individual profiles are added along the convoluted cortex, the average BOLD response was calculated as shown in Equation 6 to test the deconvolution approach. The resulting profile is shown as a thick black line in Figure 1a.

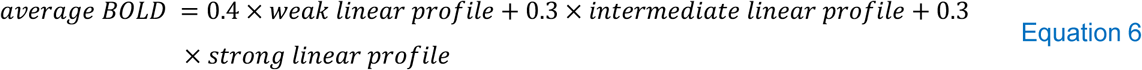

#### Multi-echo dataset

The 3D-FLASH multi-echo dataset was acquired at 7T using an isotropic voxel size of 0.75 mm (Koopmans et al., 2011). The stimulus used was a flickering checkerboard covering the central portion of the visual field. Cortical profiles were obtained by heavily sampling between corresponding vertices on the mesh generated on the WM/GM surface and on the GM/pial surface. For each pair of vertices, 100 equally spaced points were acquired within GM, and 100 equally spaced points at either side of the cortex, i.e. in the WM and pial surface. Sampling the cortex at both sides allowed for the careful realignment of cortical profiles over the activated primary visual cortex (V1) based on two landmarks, the GM/CSF boundary and the stripe of Gennari, giving profiles such as shown in Figure 1b. The magnitude and shape of the cortical BOLD profile is TE-dependent, whereas the underlying activation is the same across TEs. In the current manuscript the multi-echo data were down-sampled to 10 cortical bins within grey matter, corresponding to the cortical sampling in the cortical model.

### Assessment of the leakage-free profiles

The profiles classified as showing a weak linear profile in the ultrahigh resolution dataset have little contribution from intracortical veins. They can be considered a good approximation of leakage-free profiles, and as such, these are the profiles against which the performance of the deconvolution approach is assessed in this manuscript. For this purpose, a similarity metric between the obtained profile and the leakage free ground truth is determined using the normalised dot product (Deshpande et al., 2013), defined in Equation 7:

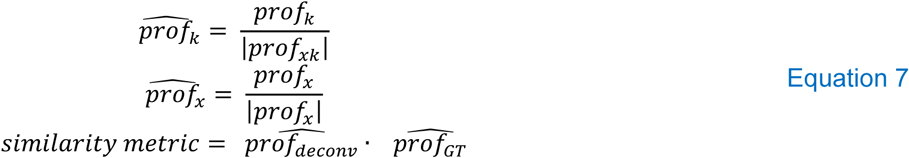

If the similarity metric is 1, the two profiles are identical or scaled versions of one another. If 0, they are orthogonal to each other.

In the present article we consider the weak linear trend profiles to represent a gold standard for the deconvolution. They are similar to spin-echo profiles, but are acquired with a normal gradient-echo technique. The weak profiles were acquired with a very high spatial resolution of 0.5 mm isotropic which may not always be attainable, or indeed desirable if the goal is to acquire data from extended regions of cortex within an acceptable acquisition time. We first consider the deconvolution of the average profiles obtained in the high resolution experiment and compare the result to the weak linear profile. We then deconvolve the profiles of the multi-echo data set and examine their similarity as a function of TE with the weak profiles. We then explore the sensitivity of the results obtained to the exact value of the p2t value chosen both for the ultrahigh resolution data set, and for the multi-echo data set. Finally, we use the multi-echo data set to determine the optimum p2t value for each subject, and averaged over subjects.

## RESULTS

### Deconvolution with the p2t on GE-BOLD activation profiles

The deconvolution results for the ultrahigh resolution dataset are shown in Figure 2. Figure 2a shows the weighted average cortical profile for each subject (one acquired twice), obtained using Equation 6, the average across all subjects is shown in black. Figure 2b shows the corresponding deconvolved profiles. The ultrahigh resolution dataset was sampled with 16 bins across the cortex, therefore, before performing the deconvolution, the p2t value from the cortical model was adjusted for the number of bins following Equation 3. These deconvolved profiles no longer show an ascending pattern towards the pial surface, and show a significantly higher similarity (p=0.00) with the corresponding, subject-specific, weak linear trend profiles than with the corresponding average cortical profiles.

The multi-echo dataset contains one activation profile per echo-time. Before applying the deconvolution for each of the echo times following Equation 2, the echo time specific p2t values were calculated using the vascular model described in (Markuerkiaga et al., 2016). The magnitude of the estimated p2t values at 7T for TEs between 4.8 and 56.1 ms are between 6 and 7.1 (see SI:Figure 2). Figure 3 shows the deconvolved profiles for each of the echo times averaged over subjects. Similar to the ultrahigh resolution dataset, the profiles no longer increase steadily from the WM/GM boundary to the GM/CSF boundary. Indeed, as shown in Figure 4, the deconvolved profiles show a significantly stronger similarity with the average of the weak linear profile in the ultrahigh-resolution dataset.

### Magnitude of the p2t

The profiles corrected for the inter-laminar leakage presented so far rely on the updated vascular model described in Markuerkiaga et al. (2016) shown in SI:Figure 2. Nonetheless, the experimental, subject specific p2t value may vary from the model-based estimation due to differences in baseline or hemodynamic conditions.

Figure 5 shows the similarity between the weak linear profile and the ultrahigh resolution dataset deconvolved with p2t values that range from 1 to 20. The similarity metric has its maximum between p2t= 5-8 and steadily decreases with increasing magnitude of the p2t (note that the similarity metric is not a very sensitive measure, so large changes in profile result in small changes in the metric.).

Figure 6 shows the effect of the magnitude of the p2t used on the deconvolved profiles by showing profiles for a range of p2t values between 1 and 20. In line with the estimation of the vascular model and the results shown in Figure 5, the curves obtained using a p2t value between 5 and 8 show flatter profiles than the rest of the p2ts and therefore stronger similarity metrics as shown in figures 5 and 6b.

**Figure 6:**
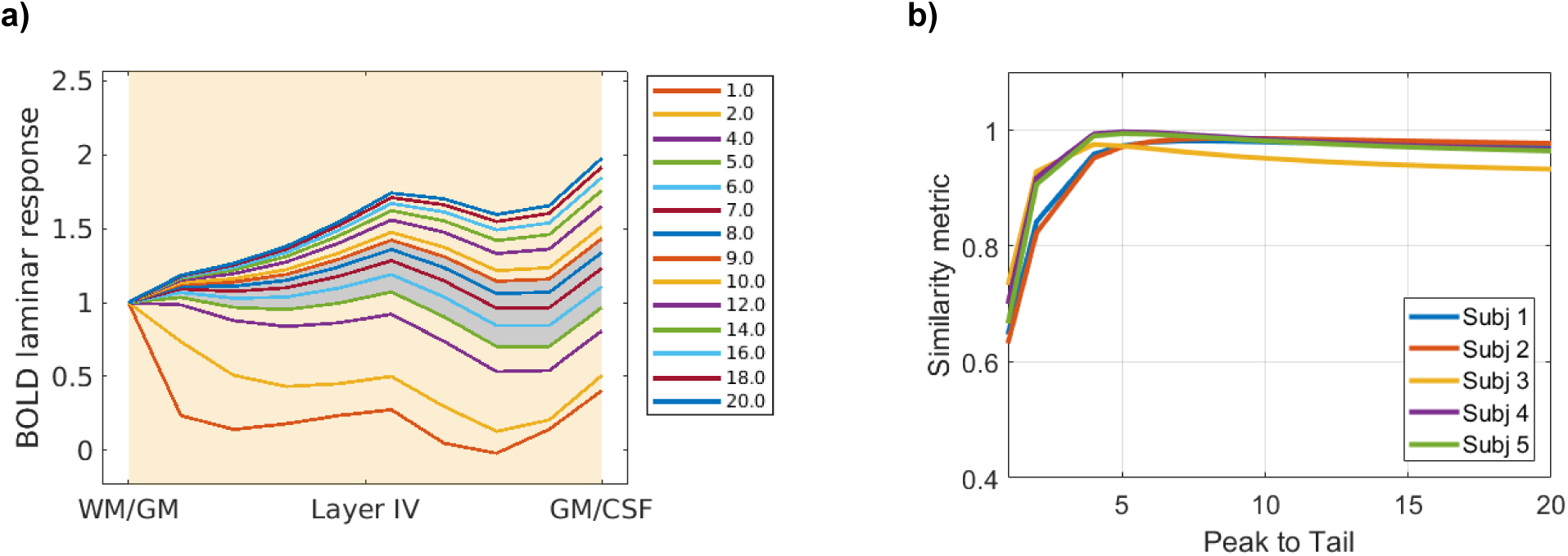
**a)** Results of deconvolving the profile at TE=T2*_GM_ averaged over all subjects in the multi-echo dataset using a range of p2t values (given in the inset). The grey-shaded region covers profiles obtained for p2t between 5 and 9, the range at which Figure 5 and Figure 6b) show a maximum. b) Similarity metric between the deconvolved multi-echo profiles at TE=T2*_GM_ with the average weak linear profile in the ultrahigh resolution dataset.

In the multi-echo dataset, measured activation profiles and p2t values are expected to be TE-dependent, but the underlying neuronal activity is not. This feature is exploited to obtain a data-based estimation of the p2t by combining the data for all echo times and solving the system in Equation 5 following a minimisation approach. Figure 7 shows the results of the p2t estimation for TE=27.6 ms, close to the T2* of grey matter at 7 T. Most of the p2t values obtained fall between 5 and 9. The results for Subject 1 represent an outlier that can be explained by the unusual shape of the profiles for this subject at longer TEs. When these longer echo times, with a poorer SNR, were not considered, then Subject 1 was no longer an outlier and the distribution of the estimated p2ts was concentrated more strongly in the range 5-9 (SI:Figure 5).

**Figure 7:**
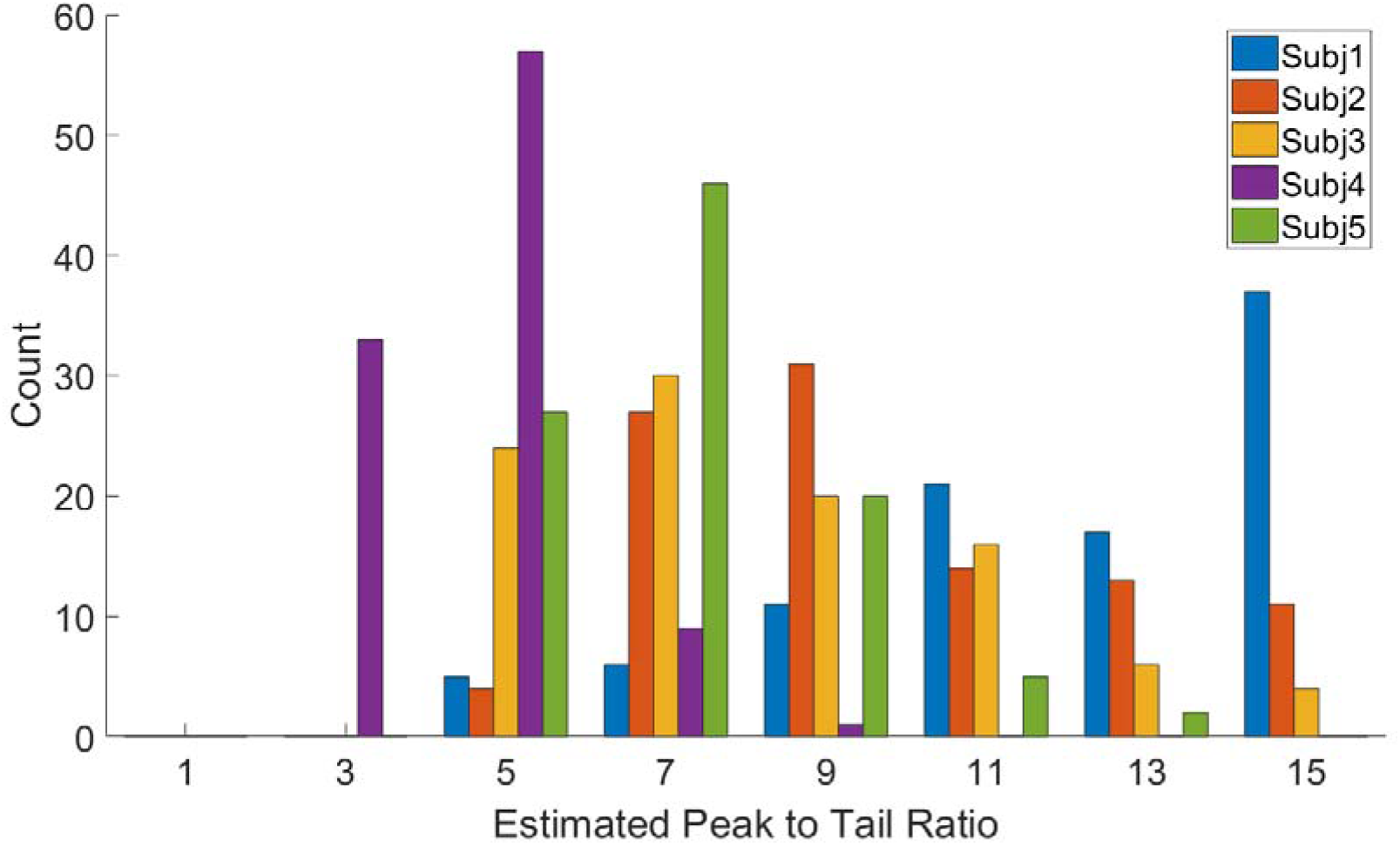
Combined histogram of the peak to tail ratio estimates at TE=27.6 ms for 100 repetitions, using data from all subjects, and estimated by applying Equation 4 to the data from the 10 echoes available. The average over subjects of the median of the p2t of the repetitions at this echo time was 7.1±2.9

## DISCUSSION

The deconvolution of GE-BOLD profiles with the physiological point spread function results in flatter cortical profiles that do not present a clear ascending pattern, in line with profiles obtained using BOLD based techniques that are less sensitive to the contribution of veins larger than 20 um in diameter, such as SE and GRASE (Jin and Kim, 2008; Kemper et al., 2015; Zhao et al., 2006). This flat profile is expected, as in both datasets the cognitive task that subjects needed to perform was to attend to a visual stimulus that consisted of a full- or semi-field flickering checkerboard. This stimulus is not expected to generate much feedback from higher order areas in the primary visual cortex (Xing et al., 2012). The deconvolution of GE-BOLD profiles with the PSF is a valid approach to eliminate the inter-laminar signal leakage through intracortical veins, while still benefiting from the higher efficiency and sensitivity achievable with GE sequences.

### Deconvolved profiles

The deconvolved profiles of the positive BOLD response show a strong resemblance to the weak linear trend profiles (see Figure 2 and SI:Figure 3). The deconvolved profiles show a significantly higher similarity degree (p=0.00) with the weak linear trend profiles than with the average measured activation profiles in that dataset. The deconvolved profiles in the multi-echo dataset also show a significantly stronger similarity with the average weak linear profiles in the high-resolution dataset, than with their corresponding measured raw profiles (p=0.027).

The deconvolved profiles in the multi-echo dataset tend to show a slight increase just above the middle of the cortex, where microvascular density shows its maximum (Duvernoy et al., 1981) and the afferent connections from the thalamus end (Rockland, 2017). This small increase is less obvious in the profiles obtained for the positive BOLD response in the high-resolution dataset from Fracasso et al. (2018), although the underlying response is expected to be similar between them. A plausible reason for this may be that in the multi-echo dataset the individual profiles were realigned using the stripe of Gennari and the GM/CSF boundary as landmarks. Slight misalignments are likely to blur the small bump and lead to a flatter profile.

### Validation of the estimated PSF and its approximation as a peak to tail ratio

Estimating the physiological point spread function is not possible/feasible in practical terms. Therefore, the deconvolution approach presented in this manuscript makes use of a model-based estimation. Using a physiological PSF obtained from an average model of cortical vasculature is expected to be a valid approximation of the subject-specific physiological PSF, as the microvascular architecture across the primary visual cortex shows little inter-subject variability (Weber et al., 2008). Similarly, the microvascular profiles between V1 and the rest of the regions in the visual cortex do not diverge considerably, although V1 shows a more pronounced peak in layer IV. The differences in micro-vascular density between regions will be accompanied by differences in the intracortical veins that drain them. The form of the PSF is not expected to vary and the fact that the parameter that describes the PSF is used as a ratio in this approach will minimise differences related to differences in the vascular density between regions.

The results in this manuscript show that a peak to tail ratio ∼6.3 for TE= 27.6 ms at 7T for 10 sampling points across the primary visual cortex adequately describes the biophysical properties of the leakage effect. If the same approach is applied to other regions then the p2t value will need to be adjusted to accommodate different cortical thicknesses using Equation 3. Various studies in laminar fMRI literature (Kok et al., 2016; Lawrence et al., 2019; Lawrence et al., 2018; Sharoh et al., 2019) divide the cortex in three layers, it that case, the equivalent peak to tail ratio to the one observed here would be 2.24.

### Combination with the spatial GLM

As mentioned in the Introduction, a BOLD profile that has been extracted using the spatial GLM still suffers from the inter-laminar leakage problem. The output of the GLM unmixing as implemented in van Mourik et al. (2019) is a signal time-course for each of the laminae. A ‘lamina’ in this context is each of the cortical bins that the cortex has been divided into for the analysis and does not have to correspond to a histological layer. The condition specific cortical activation profile obtained using this approach could further be deconvolved with the physiological PSF.

### General requirements to apply the deconvolution approach

Most importantly good segmentation and co-registration are paramount to applying the approach. If the cortical sampling deviates too much from the model due to segmentation problems, the approach will fail.

Once the cortical boundaries have been defined it is also important that the bins obtained are the same size. The model used to estimate the p2t, (Markuerkiaga et al., 2016) implicitly relies on the equivolume sampling approach (Waehnert et al., 2014). Hence, this method should be favoured over the equidistant approach, although the differences between the two have been found to be negligible for voxels with a side > 0.6 mm in the direction perpendicular to the cortical surface (Kemper et al., 2017).

As previously mentioned Equation 2 assumes that the response of the cortical vasculature to the stimulus is in a steady state and does not apply to transient cases, this will be the case for stimuli longer than 1s. Furthermore as assumed in Markuerkiaga et al. (2016) profiles should have been obtained integrating over a sufficiently large and convoluted patch of cortex that there is no preferred orientation of the venous vasculature.

### Non-BOLD sources of signal spread through layers

The Lorentzian broadening induced by the relaxation during acquisition is Δf=1/πT2* (Marques and Norris, 2018). At 7T this equates to 11.4 Hz for grey matter (T2*∼28ms) and 45.6Hz for venous blood (T2*∼7ms).

The multi-echo dataset was acquired using the 3D-FLASH sequence. Therefore, the BW per pixel in the PE direction is infinite whereas the BW/pixel in the RO direction is 240Hz/pixel. These values are above the line broadening induced by the acquisition train and hence the results presented here will not be affected.

The high-resolution dataset was acquired using 3D-EPI and the BW/pixel in the phase encoding directions was 21Hz/pixel (Fracasso et al., 2018). This is smaller than the line broadening expected for venous blood. However, the TE= 27ms was matched to T2*GM. At this echo time, the intravascular contribution to the BOLD signal is more than 10 times smaller than the extravascular contribution (Cheng et al., 2015). Hence, it can be assumed that the smearing of the profiles due to Lorentzian line-broadening is negligible. Regardless of whether deconvolution is used or not, studies focusing on laminar activation or that require high spatial accuracy should control that the Lorentzian broadening is confined to a voxel when setting the acquisition parameters.

## CONCLUSIONS

The deconvolution approach has been applied to GE-BOLD profiles obtained at 7T acquired using similar stimuli but different sequences, resolutions and processed with different number of samples across the cortex. The deconvolved profiles no longer show an ascending shape, in-line with the profiles obtained without a strong contribution from large vessels. The deconvolution approach offers a means to obtain inter-laminar leakage-free profiles, while still benefitting from the higher efficiency and sensitivity of GE-BOLD based functional acquisition methods.

## Supporting information

SI for laminar deconvolution

## ACKNOWLEDGEMENTS

This work was supported by the Initial Training Network, HiMR, funded by the FP7 Marie Curie Actions of the European Commission (FP7-PEOPLE-2012-ITN-316716). The authors would like to thank Peter Koopmans, Alession FracassoF, Natalia Petriodu and their collaborators for providing high resolution GE-BOLD cortical profiles and Eli Alberdi for critical feedback on the minimization approach. Part of this work has been presented in the 24th ISMRM Annual Meeting in Singapore.

