## Supplementary material for "Estimation of Laminar BOLD Activation Profiles using Deconvolution with a Physiological Point Spread Function": SI for laminar deconvolution

### SI part 1

In this manuscript, an approximation of the PSF estimated in Markuerkiaga et al. (2016) has been used. Namely, the continuous laminar PSF has been approximated as a peak in the layer of activation followed by a constant peak in the layers downstream. This PSF can be described by a single parameter, the peak to tail ratio (p2t). In addition, the same PSF for all layers (albeit with different tail lengths) has been considered across the cortex. Here, we will show the effect that these approximations have on the deconvolved profiles.

SI:Figure 1 shows the results obtained using the model developed in (Markuerkiaga et al., 2016). In this example, the same metabolic and hemodynamic response was simulated in all layers, i.e. $\Delta SO_2$ and $\Delta CBV$ [%] were the same in all layers. The dark grey curve shows the simulated GE-BOLD activation profile obtained, whereas the light grey curve shows the simulated SE-BOLD activation profile. Note, that the SE profile is not completely flat, this is especially clear around 0.4 of the cortical depth, where the microvascular density is the highest.

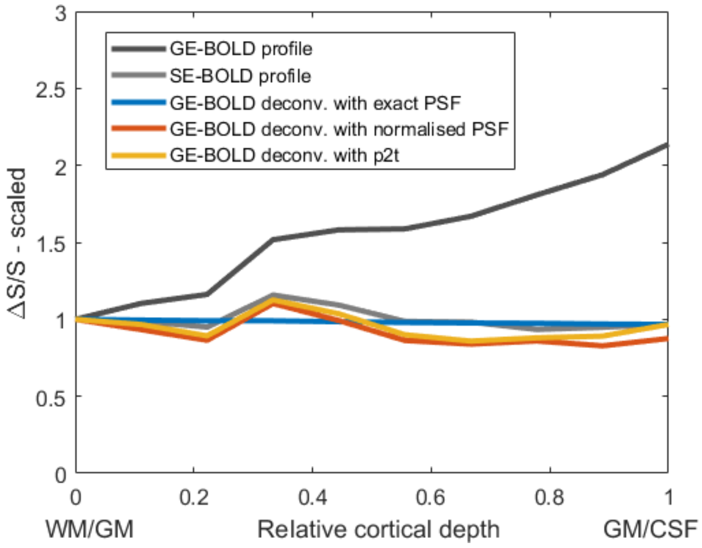


**SI:Figure 1**: Simulated profiles obtained using the vascular model and PSFs presented in (Markuerkiaga et al., 2016). All profiles have been normalised to their values at the WM/GM boundary to facilitate comparison. The Gradient Echo (dark grey) profile shows an ascending pattern, whereas the Spin Echo profile (light grey) follows the microvascular density across the cortex (see Figure 1 in Markuerkiaga et al. (2016) for the microvascular density in the model). No noise was added to the simulated GE-BOLD profile. The profiles in colour show the results obtained by deconvolving the GE-BOLD profile with three versions of the physiological point spread function: i) If the exact PSF shown in Figure 5b in Markuerkiaga et al. (2016) is used, then an activation profile normalised with the local vascular density is obtained (in blue). ii) If the normalised PSF is used, i.e. each depth specific PSF normalised by its value at the layer of activation, a profile with some microvascular weighting is obtained (red profile). iii) If the tail of the PSF is approximated to be flat and the PSF is characterised by a single p2t values across the cortex (as in the present article), a profile with some microvascular weighting is still obtained (yellow profile).

The use of the PSF and the effect of the simplifications made are discussed next. The absolute values of the PSFs s shown in Figure 5b of (Markuerkiaga et al., 2016) differ. This is mainly due to differences in the microvascular density across the cortex. If deconvolution is performed using the exact PSF values, laminar BOLD activation profiles normalised by the underlying microvascular density will be obtained (blue profile in SI:Figure 1, note that the same strength of hemodynamic response was simulated). This profile is flat and does not present the small protuberance found in the SE profile. This reflects the assumption of an initial flat response across the layers.

If normalised PSFs are used (but the PSF is still laminar specific) then a curve is similar to the SE curve (orange curve in SI:Figure 1) is obtained (orange curve). If, as done in this manuscript, the tail is considered to be flat, i.e. 1 in the laminar peak and a constant 1/p2t in the tail, then the yellow curve in SI:Figure 1 is obtained, which is similar to the case in which the normalised PSF is used (compare yellow and orange curves in SI:Figure 1).

The main difference between using the normalised PSF (yellow and orange curves in SI:Figure 1) and non-normalised PSF (blue curve in SI:Figure 1) is that the normalised versions show variations in the profile due to differences in the microvascular density across the cortex, similar to the SE profile. In other words, although this approach compensates for the leakage through intracortical veins, the differences in the BOLD response due to differences in the laminar microvascular density are still present.

#### Part 2

##### Modification of the model parameters to accommodate a longer venous T2* at 7T

Following (Uludağ et al., 2009), the p2t estimated in (Markuerkiaga et al., 2016) assumed that intravascular contribution to the BOLD signal was negligible at 7T and beyond. Experimental BOLD fMRI measurements at 7T have shown that the intravascular contribution is not completely vanished. In order to apply the deconvolution with the estimated p2t on the ultra-high resolution and multi-echo datasets in this manuscript, the model parameters in (Markuerkiaga et al., 2016) have been modified to simulate profiles with some intravascular contribution and obtain a more accurate p2t estimate.

The choice of the parameters was empirically determined so that the intravascular contribution to the BOLD signal was ~8%. In the modelling paper relaxation times, $R2^{(*)}$ , is the sum of a relaxation time that is independent of the dHb concentration, $R2^{(*)}_0$, and one depends on the dHb concentration $R2^{(*)}_{dHb}$.

| **No intravascular contribution** (Markuerkiaga et al., 2016) | **With intravascular contribution** (This manuscript) |
|---|---|
| 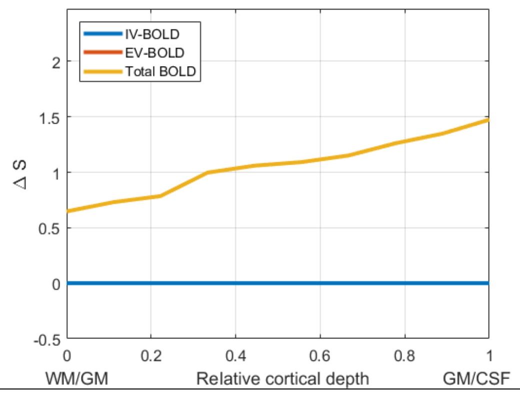  | 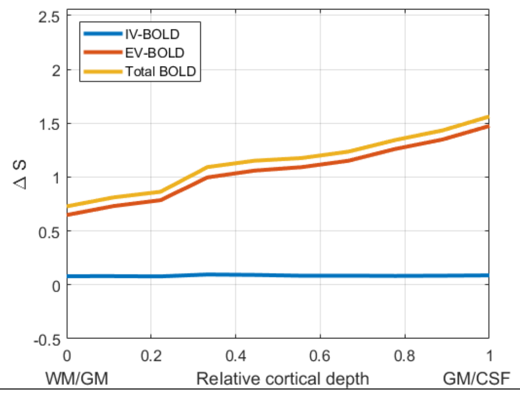  |
| Intravascular relaxation time:<br>$R2^*_{in} = \infty$ | $R2^*_{0\_in} = R2_{0\_in} *4$<br>$R2^*_{dHb\_in} = R2_{\_dHb}$<br>Intravascular relaxation time:<br>$R2^*_{in} = R2^*_{0\_in} + R2^*_{dHb\_in}$ |

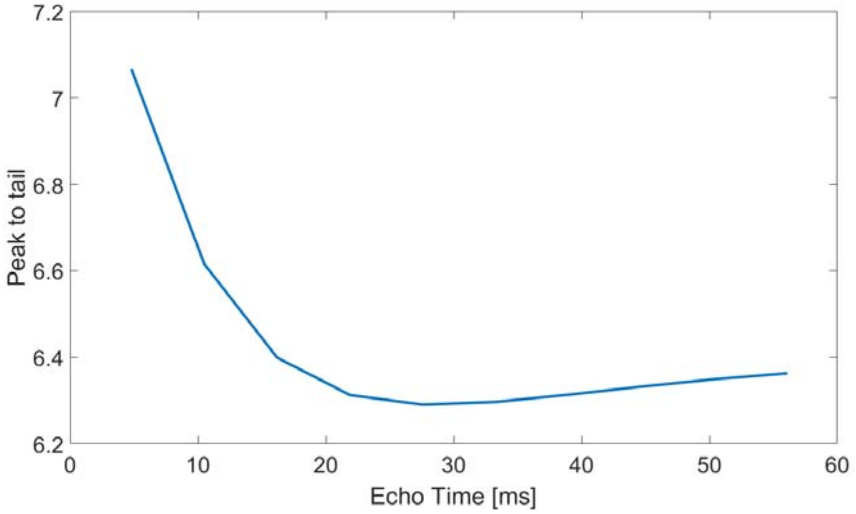


**SI:Figure 2**: Magnitude of the peak to tail ratio at the echo times used in the multi-echo dataset for 7T. Calculated using the simulation of Markuerkiaga et al. (2016), but with the modified intravascular relaxation time so that the IV contribution to the BOLD signal has not completely vanished at TE=$T2^*_{GM}$ but is ~ 8%.

**Part 3. Estimation of the p2t adjustment for arbitrary number of bins**

Following Equation 1 in the body of the manuscript, the BOLD response at a given cortical bin is obtained by the following expression:

$$y_i = b_i * peak + \sum_{k=1}^{i-1} b_k * tail$$

The total response across the cortex, *A*, is obtained by summing over all bins.

$$A = \sum_i y_i \Delta x = \sum_i \left( b_i * peak + \sum_{k=1}^{i-1} b_k * tail \right) \Delta x$$

where $\Delta x$ is the thickness of the bin. For the p2t adjustment we seek, without loss of generality, we can solve the system for the specific case in which $b_i = 1, \forall i$. In this case, the expression is simplified to:

$$A = \sum_{i=1}^{n} \left( peak + \sum_{k=1}^{i-1} tail \right) \Delta x$$

As $p2t = \frac{peak}{tail}$ and $\Delta x = \frac{corticalThickness}{n}$, where n is the number of bins across the cortex:

$$A = \sum_{i=1}^{n} \left( p2t + \sum_{k=1}^{i-1} 1 \right) tail \, \frac{corticalThickness}{n}$$

$$A = \sum_{i=1}^{n} (p2t + i - 1) * tail * \frac{corticalThickness}{n} = \left( n * (p2t - 1) + \sum_{i=1}^{n} i \right) * tail * \frac{corticalThickness}{n}$$

As $\sum_{i=1}^{n} i = \frac{n(n+1)}{2}$ :

$$A = \left( p2t - 1 + \frac{(n+1)}{2} \right) * tail * corticalThickness = \left( \frac{2 * p2t + n - 1}{2} \right) * tail * corticalThickness$$

The p2t depends on the number of bins, *n*, used but the total BOLD response should be independent of the number of bins. Therefore,

$$A(n_1) = A(n_2)$$

$$\left( \frac{2 * p2t_{n_1} + n_1 - 1}{2} \right) * tail_{n_1} * corticalThickness = \left( \frac{2 * p2t_2 + n_2 - 1}{2} \right) * tail_{n_2} * corticalThickness$$

$$\left(2 * p2t_{n_1} + n_1 - 1\right) * tail_{n_1} = (2 * p2t_2 + n_2 - 1) * tail_{n_2}$$

$$p2t_{n_1} = \left(\frac{tail_{n_2}}{tail_{n_1}}(2 * p2t_{n2} + n_2 - 1) + 1 - n_1\right) * \frac{1}{2}$$

The magnitude of the tail is proportional to the volume of the unit element that is being drained. This volume is inversely proportional to the number of bins used. Hence, $\frac{tail_{n_2}}{tail_{n_1}} = \frac{n_1}{n_2}$

$$p2t_{n_1} = \left(\frac{n_1}{n_2}(2 * p2t_{n2} + n_2 - 1) + 1 - n_1\right) * \frac{1}{2}$$

$$p2t_{n_1} = p2t_{n2}\frac{n_1}{n_2} + \frac{(n_2 - n_1)}{2n_2}$$

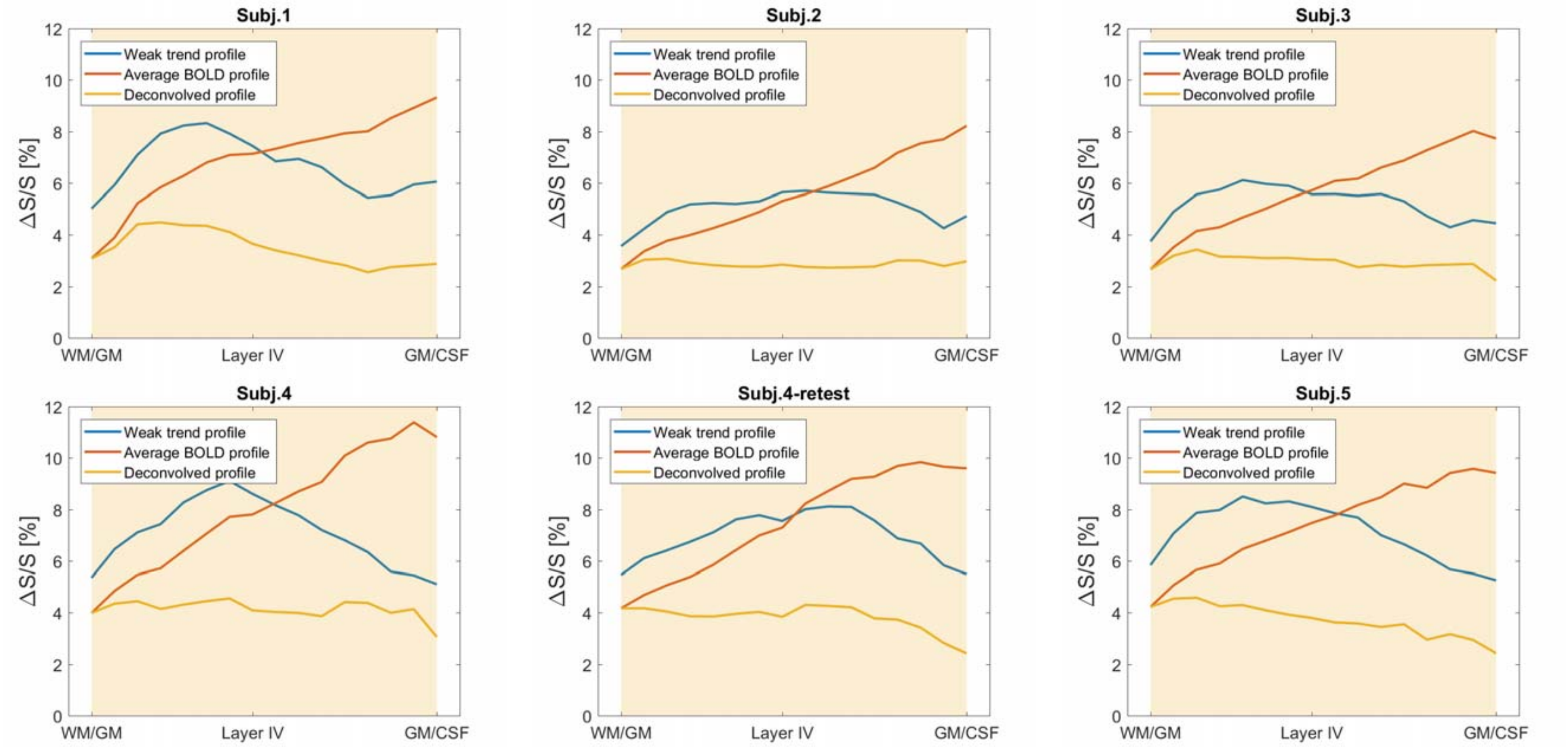


**SI:Figure 3:** Single subject profiles obtained with the ultrahigh-resolution data for each of the 6 measurements (5 subjects, one subject measured twice) for the positive BOLD response. The weakly linear profile (blue), the average of the weak, middle and strong linear profiles calculated as explained in the manuscript (red) and the average profile deconvolved with the estimated peak to tail ratio (yellow).

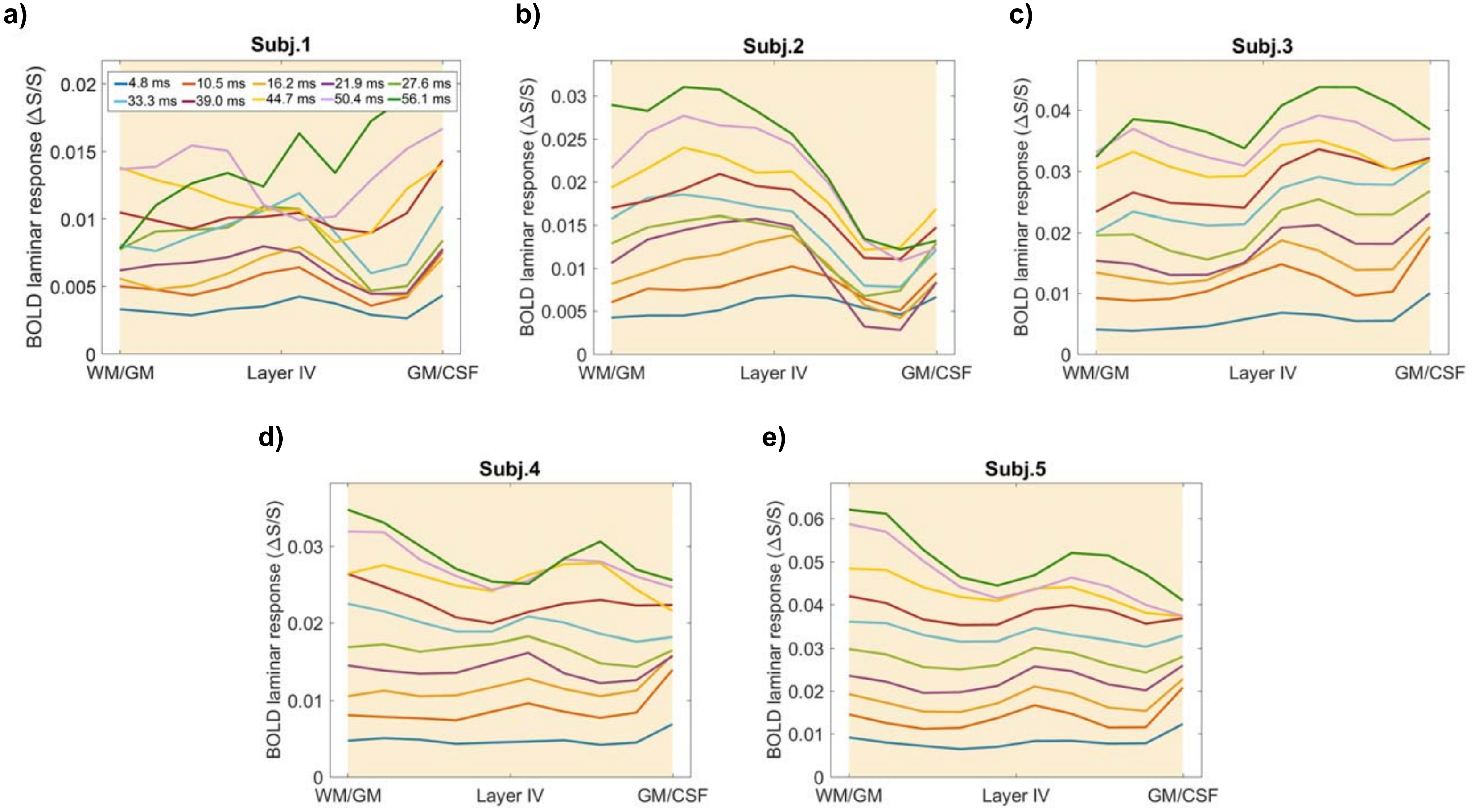


**SI:Figure 4:** Deconvolved laminar BOLD responses per subject obtained for the multi-echo dataset applying Equation 1 using the corresponding p2t from the model for two commonly used TEs, graphs **a)** to **e)**. The difference in the magnitude of the profiles between subjects reflects the difference in the average response between subjects.

As expected, the profiles obtained for different TEs differ in the average magnitude of the response, but their shape is very similar.

The deconvolved profiles shown in SI:Figure 4 are the underlying responses, β, obtained by applying SI Equation 1 given below. The TE-specific p2t values obtained from the model in (Markuerkiaga et al., 2016) have been used.

$$\begin{bmatrix} y_1 \\ y_2 \\ \vdots \\ y_n \end{bmatrix} = \begin{bmatrix} 1 & 0 & \dots & 0 \\ 1/p2t & 1 & & 0 \\ 1/p2t & 1/p2t & \ddots & 0 \\ 1/p2t & 1/p2t & \dots & 1 \end{bmatrix} \begin{bmatrix} \beta_1 \\ \beta_2 \\ \vdots \\ \beta_n \end{bmatrix} + \begin{bmatrix} \varepsilon_1 \\ \varepsilon_2 \\ \vdots \\ \varepsilon_n \end{bmatrix} \qquad \text{SI:Equation 1}$$

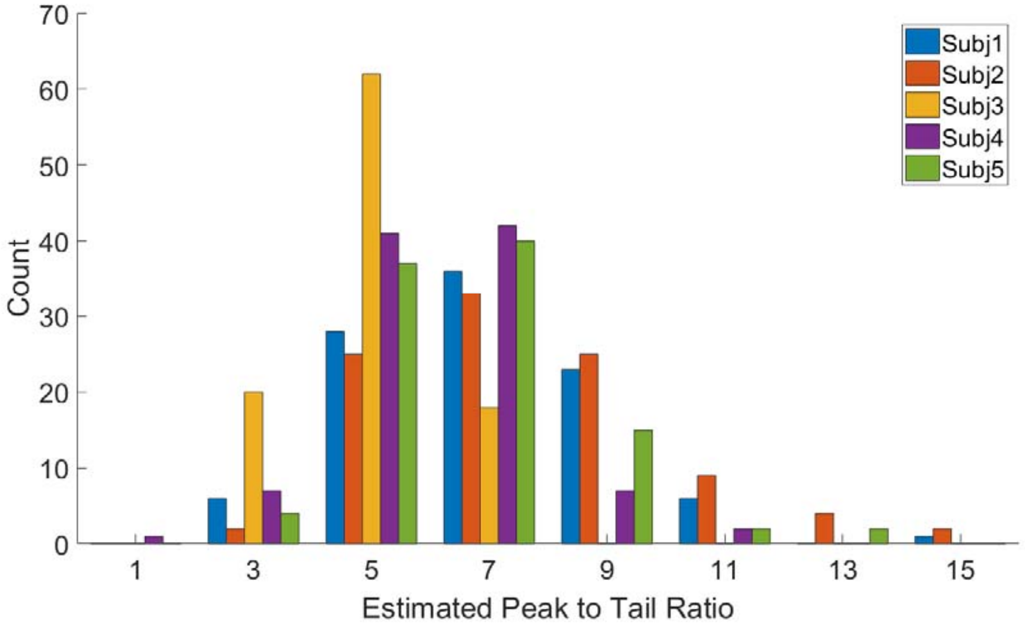


**SI:Figure 5: a)** Combined histogram of the peak to tail ratio estimations for 100 repetitions of all subjects for TE=27.6 ms obtained applying Equation 4 using only two echoes, TE= 21.9 ms and TE=27.6 ms. The average over subjects of the median of the p2t of the repetitions was 6.6±0.9
